# Efficient *hyperactive piggyBac* transgenesis in *Plodia* pantry moths

**DOI:** 10.1101/2022.10.19.512940

**Authors:** Christa Heryanto, Anyi Mazo-Vargas, Arnaud Martin

## Abstract

While *piggyBac* transposon-based transgenesis is widely used in various emerging model organisms, its relatively low transposition rate in butterflies and moths has hindered its use for routine genetic transformation in Lepidoptera. Here, we tested the suitability of a codon-optimized *hyperactive piggyBac* transposase (*hyPBase*) in mRNA form to deliver and integrate transgenic cassettes into the genome of the pantry moth *Plodia interpunctella*. Co-injection of *hyPBase* mRNA with donor plasmids successfully integrated 1.5-4.4 kb expression cassettes driving the fluorescent markers EGFP, DsRed, or EYFP in eyes and glia with the *3xP3* promoter. Somatic integration and expression of the transgene in the G_0_ injected generation was detectable from 72-hr embryos and onward in larvae, pupae and adults carrying a recessive white-eyed mutation. Overall, 2.5% of injected eggs survived into transgene-bearing adults with mosaic fluorescence. Subsequent outcrossing of fluorescent G_0_ founders transmitted single-insertion copies of *3xP3::EGFP* and *3xP3::EYFP* and generated stable isogenic lines. Random in-crossing of a small cohort of G_0_ founders expressing *3xP3::DsRed* yielded a stable transgenic line segregating for more than one transgene insertion site. We discuss how *hyPBase* can be used to generate stable transgenic resources in *Plodia* and other moths.

## INTRODUCTION

Lepidoptera is a large insect order that comprises 160,000 species (Kristensen et al., 2007; Roskov et al., 2013), including a wide range of agricultural pests and ecosystem service providers, as well as important model systems for research in conservation biology, ecology, and evolutionary biology. In order to foster the potential of lepidopteran insects for functional genetics beyond the silkworm flagship system, for which transgenic resources already exist, we are developing the pantry moth *Plodia interpunctella* (hereafter *Plodia*; abbr. *Pi*), or Indianmeal moth, as an alternative laboratory organism amenable to routine genome editing and transgenesis. *Plodia* is a worldwide pest of stored food products, and exhibits convenient laboratory features that make it a promising system for the long-term maintenance of isogenic lines. In addition to its relatively short life cycle (25 days at 28°C) and ease of culture on a low-cost diet (Silhacek and Miller, 1972), *Plodia* cultures are resilient to inbreeding (Bartlett et al., 2018). Mass egg-laying can be stimulated by exposing their highly fecund females (Mbata, 1985) to CO_2_ gas, a property that allows the collection of synchronized embryos within the time frame of the first cell divisions, thus facilitating genetic transformation by microinjection (Dyby and Silhacek, 1997; Bossin et al., 2007). Finally, several genome assemblies and several transcriptomic resources have been published in this species (Harrison et al., 2012; Tang et al., 2017; Roberts et al., 2020; Heryanto et al., 2022; Kawahara et al., 2022).

Transgenesis techniques based on the *piggyBac* transposase (*PBase*) have been successfully implemented in a wide variety of insect model organisms and beyond (Handler, 2002; Gregory et al., 2016; Laptev et al., 2017). Butterflies and moths were shown to have transposition rates an order of magnitude lower than in beetles, mosquitoes and flies (Gregory et al., 2016), making routine transgenesis more challenging in the Lepidoptera order. A modified version of the transposase dubbed *hyperactive piggyBac* (*hyPBase*) was isolated from a mutant screen in 2011 (Yusa et al., 2011). *hyPBase* was later shown to dramatically increase transformation rates in flies and honeybees compared to its native version (Eckermann et al., 2018; Otte et al., 2018), and was also shown to provide practical transformation rates in *Spodoptera* noctuid moths (Chen and Palli, 2021).

Previously, delivery of the original *PBase* as a helper plasmid into *Plodia* syncytial embryos resulted in somatic transformation of fluorescent markers, but its efficiency for germline transformation was not reported (Bossin et al., 2007). Here, we extend the assessment of *hyPBase* transgenesis in Lepidoptera with a focus on the pyralid moth *P. interpunctella*, a pest of stored foods that is amenable to genome editing and genetic transformation (Bossin et al., 2007; Heryanto et al., 2022). In the current study, we injected an insect codon-optimized *hyPBase* as a mRNA (Otte et al., 2018) and monitored both the somatic and germline transformation rates of fluorescent markers driven by the *3xP3* promoter, a canonical promoter with strong activity in the ocular and glial tissues in Lepidoptera and other insects (Berghammer et al., 1999; Horn et al., 2002; Thomas et al., 2002). This approach robustly generated transgenic lines carrying various fluorescent protein markers, illustrating the suitability of *hyPBase* for routine genetic transformation in *Plodia* pantry moths. We discuss future strategies for establishing transgenic lines in emerging laboratory systems for lepidopteran functional genomics.

## RESULTS

We tested the suitability of *hyPBase* for transgenesis, using three donor plasmids that drive the expression of the fluorescent markers EGFP, DsRed, and EYFP. For each experiment, we report the levels of somatic transformation observed in the G_0_ injected generation, as well as our observations carrying the transgenes into further G_1-3_ generations.

### hyPBase delivery of a 4.4 kb insert expressing 3xP3::EGFP

A practical transgenesis method must allow the delivery of relatively large cargos of several kilobases. To test the efficiency of *hyPBase*, we generated a *piggyBac* donor plasmid with a 4.4 kb insert with both a transgene and a transgenesis marker (Fig. 1B). The cassette consisted of the *mScarlet* red fluorescent protein flanked by 3’ and 5’UTR regions of the *nanos-O* gene *Plodia* homolog, a germline determinant selected on its apparent specificity to gonadic tissues (Nakao and Takasu, 2019; Xu et al., 2022). As a transgenesis marker, we used a *3xP3::EGFP* marker that labeled ocular tissues during previous somatic *piggyBac* transformation attempts in *Plodia* (Bossin et al., 2007). First, we injected this plasmid without *hyPBase* mRNA to control for episomal expression of the *3xP3::EGFP* driver. These injections showed strong EGFP expression in large internal cells 48 h post-injection, suggesting episomal expression from the embryo vitellophages (Fig 2A). However, this signal was lost in 72-h old embryos, which only showed background levels of fluorescence or external autofluorescence artifacts at injection sites (Fig 2B). Thus, episomal expression of injected plasmids dissipates by 72 h of embryonic development and should not interfere with the screening of successful integration events at this stage and onwards.

**FIGURE 1.**
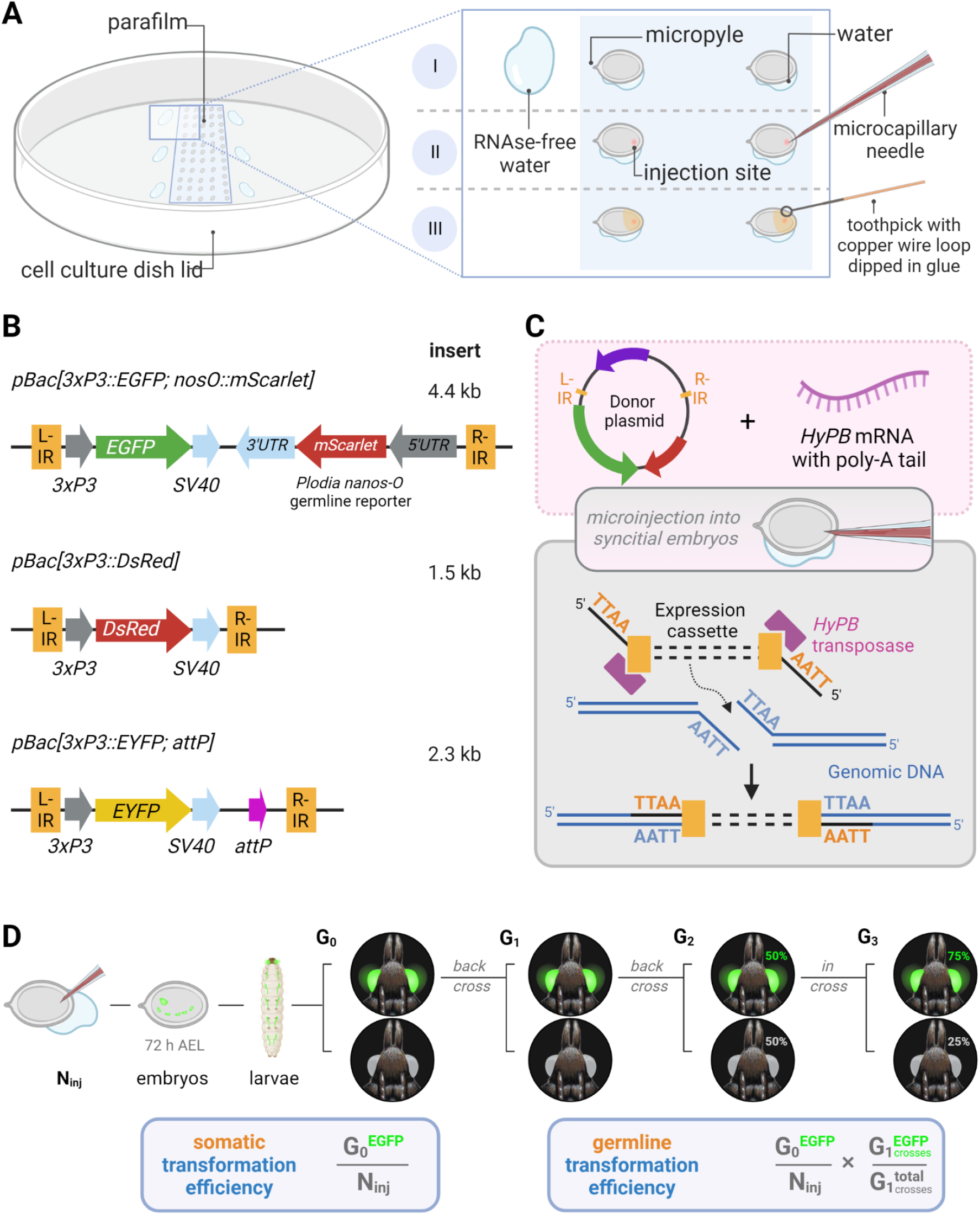
Microinjection procedure and transgenic constructs for the testing of *hyperactive piggyBac* transformation in *Plodia*. **(A)** Microinjection of *P. interpunctella* syncytial embryos. Gravid females oviposit *en masse* after CO_2_ narcosis, and eggs are collected and oriented on a parafilm strip in a tissue culture dish. A wet brush is used to position eggs, with water contact helping firm adhesion to the parafilm (I). Microinjection is performed on the side opposite to the micropyle (II). Peripheral droplets of water are used to periodically flush the injection capillary of yolk. Eggs are sealed with glue following injection (III). **(B)** Expression cassettes of donor plasmids carrying *3xP3* eye and glia fluorescent markers. IR = *piggyBac* internal repeats (L, left ; R, right). **(C)** Transposon-mediated random integration following the injection of donor plasmid and *hyPBase* mRNA. **(D)** Somatic transformation efficiency (%) is equivalent to the number of potential G_0_ founders obtained out of 100 injected eggs. Germline transformation efficiency (%) factors proportion of transgenic G_1_ broods obtained from G_0_ outcrosses. N_inj_ = number of injected eggs. Made with Biorender.

**FIGURE 2.**
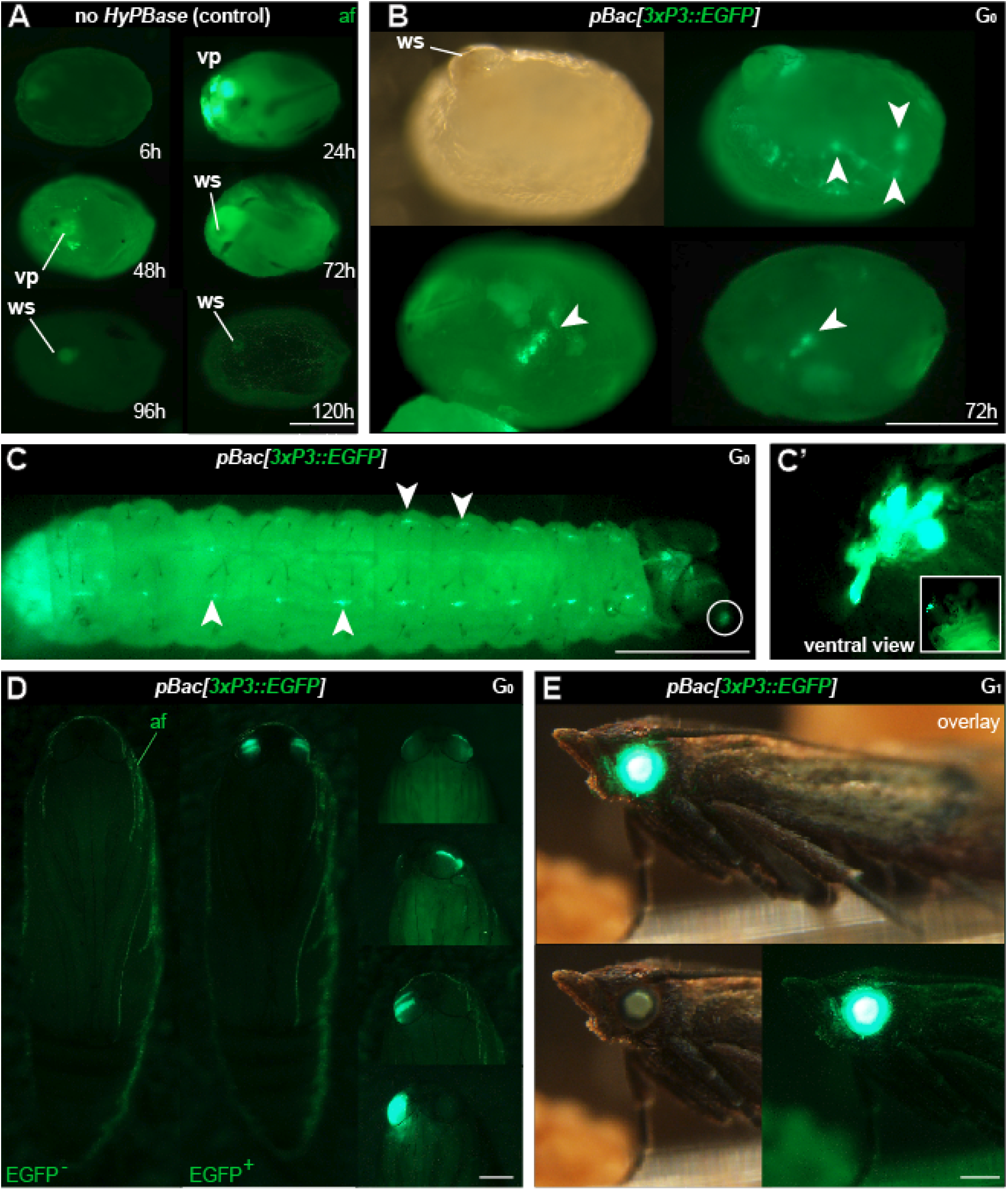
Phenotype of transgenic *Plodia* expressing EGFP in eyes and putative glia. **(A)** Control injections of *pBac[3xP3::EGFP]* show variable levels of green autofluorescence (af), most markedly at the injection wound site (ws). Episomal expression of EGFP in vitellophages (vp) is intense 24 h post-injection, reduced to background level after 48 h. **(B)** Donor *pBac[3xP3::EGFP] + hyPBase* mRNA injections resulted in *3xP3::EGFP* expression, emerging as nervous system markings around 72 h post-injection (arrowheads). 23.7% of injected G_0_ eggs (262/1104) showed a similar fluorescence during screening. **(C)** *3xP3::EGFP* expression in a first instar larva, in ganglia of the Central Nervous System (consistent with an expected glial reporter activity of *3xP3*), and in ocellar stemmata (circled, magnified in C’). **(D)** G_0_ mosaics of *3xP3::EGFP* expression in pupal eyes. An EGFP-negative pupa is shown on the left for reference. **(E)** *3xP3::EGFP* expression in a G_1_ *Plodia* adult with non-mosaic expression of EGFP in the eye (bottom right). EGFP is also visible in the brightfield (bottom right), with a green tint of the compound eyes in the *Pi_wFog* recessive white-eyed strain. Scale bars: A-C’ = 200 μm; D-E = 500 μm.

We then co-injected the donor plasmid *pBac[3xP3::EGFP; nosO::mScarlet]* with a *hyPBase* mRNA and monitored somatic transformation efficiencies throughout the G_0_ generation. In order to facilitate the screening of fluorescence, all experiments were performed in the *Pi_wFog* white-eyed strain that is devoid of screening pigments in eye tissues and also shows increased larval translucency (Heryanto et al., 2022).Transformed embryos and first instar hatchlings showed ocellar and glial EGFP fluorescence (Fig. 2C-D), with 23.7% of injected eggs showing EGFP in 72-h embryos (Table 1). Injections produced viable larvae with persistent ocellar fluorescence, as well as eye fluorescence in pupae and adults (Fig. 2E-F). Over several replicated experiments, we found that 16% of injected eggs resulted in pupae, of which 18.6% were EGFP^+^. Taking into account occasional pupal failure observed in normal rearing conditions, we determined that 2.5-3% of injected eggs become viable and fertile G_0_ somatic transformants.

**TABLE 1.**
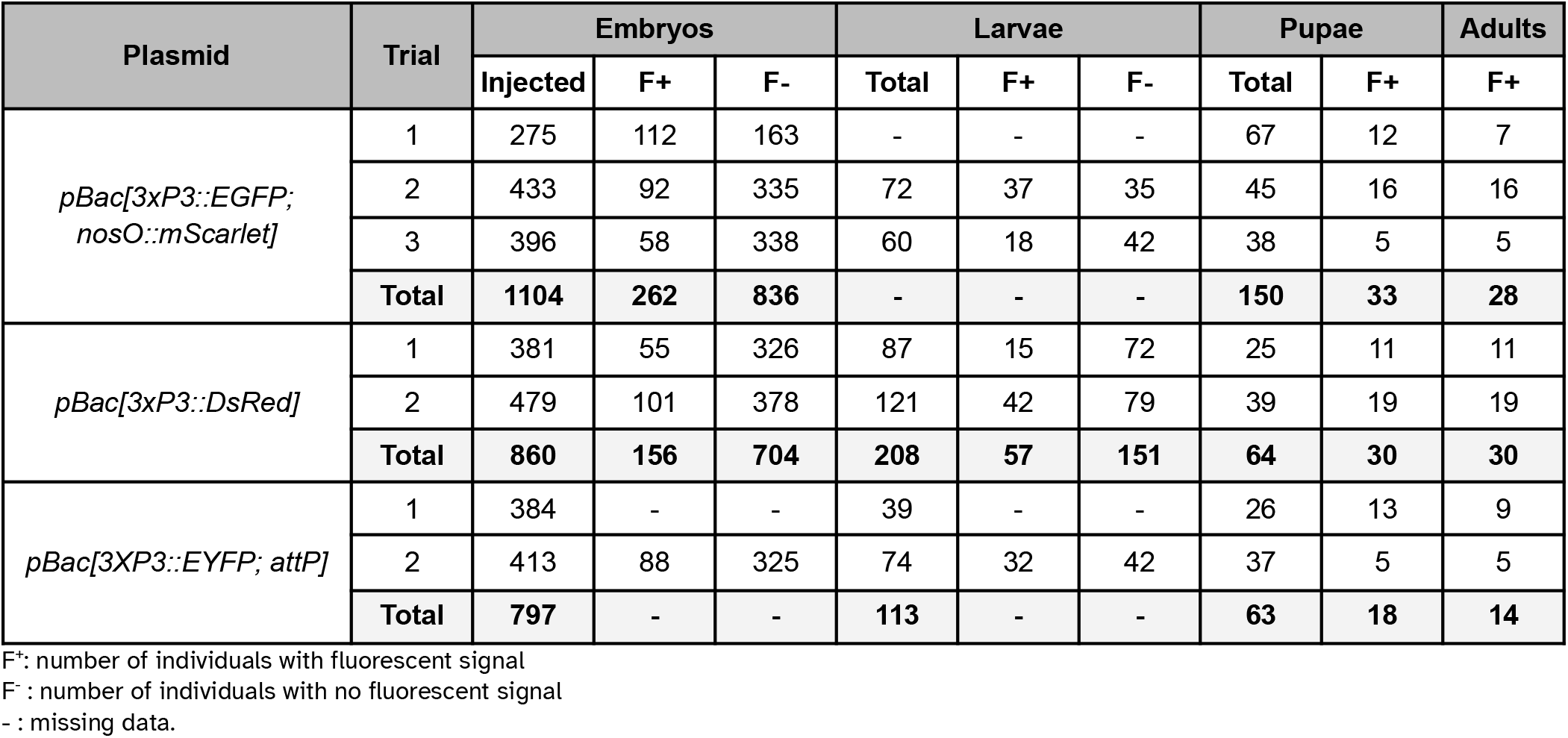
| G_0_ phenotypes of *Plodia* injected with *pBac* donor plasmids and *hyPBase* transposase mRNA.

Next, we tested germline transmission by back-crossing G_0_ EGFP^+^ individuals to uninjected stock (Table 2). Out of 6 fertile pairs, 50% yielded EGFP^+^ G_1_ progeny, suggesting a practical level of germline mobilization among G_0_ founders. This result is mitigated by the fact that only 6 out a total of 16 single-pair matings (37.5%) generated in our conditions. This establishes a germline efficiency rate of 0.94% (Fig. 1D, GTE = 6/16 × 2.5% G_0_ founders), meaning that for 1000 G_0_ embryos injected, 9.4 embryos will survive as fertile founders passing the transgene to the G_1_ generation.

**TABLE 2.** | Subsequent crossing of transgenic *Plodia* G_0_ founders.

| G<sub>1</sub> experiments | No. of G<sub>0</sub> crosses + strategy |  | No. of fertile G<sub>1</sub> broods |  | Crossing success rate |
| --- | --- | --- | --- | --- | --- |
|  |  |  | with F<sup>+</sup> | no F<sup>+</sup> |  |
| *pBac[3xP3::EGFP; nosO::mScarlet]* | 16 | BC to wFog | 3 | 3 | 37.5% (6/16) |
| *pBac[3xP3::DsRed]* | 18 | BC to wFog | 0 | 2 | 11.1% (2/18) |
|  | 1-4 | G0 in-cross | 1 | NA | container of 5 G<sub>0</sub>s (F<sup>+</sup>) |
| *pBac[3XP3::EYFP; attP]* | 14 | BC to wFog | 1 | 6 | 50% (7/14) |
| G<sub>2</sub> experiments | No. of G<sub>1</sub> fertile crosses + strategy |  | Total no. of G<sub>2</sub> progeny |  | Expected ratio if G<sub>1</sub> heterozygous at single-insert |
|  |  |  | F<sup>+</sup> pupae | F<sup>-</sup> pupae |  |
| *pBac[3xP3::EGFP; nosO::mScarlet]* | 3 | BC to wFog | 184 | 189 | yes (1:1) |
| *pBac[3xP3::DsRed]* | 3 | BC to wFog | 167 | 92 | no |
| *pBac[3XP3::EYFP; attP]* | 2 | BC to wFog | 101 | 87 | yes (1:1) |
|  |  | G1 in-cross | 55 | 23 | yes (3:1) |
| G<sub>3</sub> experiments | No. of G<sub>2</sub> fertile crosses + strategy |  | Total no. of G<sub>3</sub> progeny |  | Expected ratio if G<sub>2</sub> heterozygous at single-insert |
|  |  |  | F<sup>+</sup> pupae | F<sup>-</sup> pupae |  |
| *pBac[3xP3::EGFP; nosO::mScarlet]* | 5 | G2 in-cross | 102 | 34 | yes (3:1) |
| *pBac[3xP3::DsRed]* | mixed | G2 in-cross | 189 | 58 | yes (3:1) |
| *pBac[3XP3::EYFP; attP]* | mixed | G2 in-cross | 189 | 51 | yes (3:1) |

F<sup>+</sup>: number of individuals with fluorescent signalF<sup>-</sup>: number of individuals with no fluorescent signal

As we wanted to assess whether *hyPBase* would allow the rapid isolation of single-insertion lines, we needed to test if transgenes were integrated into multiple copies per G_0_ gamete, or if they could cause sterility. EGFP^+^ G_1_ individuals (N= 3) were back-crossed (Table 2) and produced a mean of 61 EGFP^+^ adults out of 124 emerged G_2_ per cross (49.3%), showing no statistical difference from an expected 50% ratio of a single insertion event (0.06< ^*2*^<0.46; *df* =1; 0.10< *p*< 0.80). Likewise, a total of five subsequent in-crosses (G_2_ EGFP^+^ x G_2_ EGFP^+^) each resulted in positive offspring ratios close to the expected 75% (0.06< ^*2*^<0.44; *df* =1; 0.507< *p*< 0.80). Of note, the *Plodia-nosO::mScarlet* transgene failed to drive red fluorescent signals detectable by epifluorescent and confocal microscopy in dissected ovaries, and we will explore the activity of alternative germline-driving promoters in the future (Nakao and Takasu, 2019; Xu et al., 2022). Overall, these data demonstrate that *hyPBase* provides practical transformation rates for a relatively large cargo insert, with at least 2.5% of injected zygotes yielding potential founders ready for isogenic line establishment after only one or two generations of backcrossing.

### Evaluation of a 3xP3::DsRed donor vector

To expand the toolkit of transgenesis markers, we sought to test the activity of a *pBac[3xP3::DsRed]* donor vector for the screening of red eye fluorescence. We used the *pHD-DsRed* plasmid available through Addgene (Gratz et al., 2014, 2015), which carries a 1,146 bp *3xP3::DsRed-SV40* cassette tightly flanked by *piggyBac* internal repeats. Control injections without *hyPBase* mRNA revealed weak episomal expression in vitellophages and red background fluorescence (Fig. 3A). Injection sites, which show nonspecific autofluorescence under EGFP filter sets, do not fluoresce in the red channel. *hyPBase*-mediated insertion of *pBac[3xP3::DsRed]* resulted in glial signals in 14.4% of injected 72-h AEL (after egg-laying) embryos, but intriguingly, no signal in the head region. Likewise, larval transformants showed sporadic signals in abdominal regions, seemingly nervous ganglia, but these patterns were always mosaic (Fig. 3C). About 25/30 G_0_ DsRed^+^ pupae (83%) exhibited DsRed expression in the body (Fig. 3D, G_0_ DsRed^body^). DsRed fluorescence in the head region was observed in only 5 G_0_ pupae (Fig. 3D, G_0_ DsRed^eye^), but its expression failed to reproduce the *3xP3::EGFP* signal pattern in ocelli and eye tissues (Fig. 2D).

**FIGURE 3.**
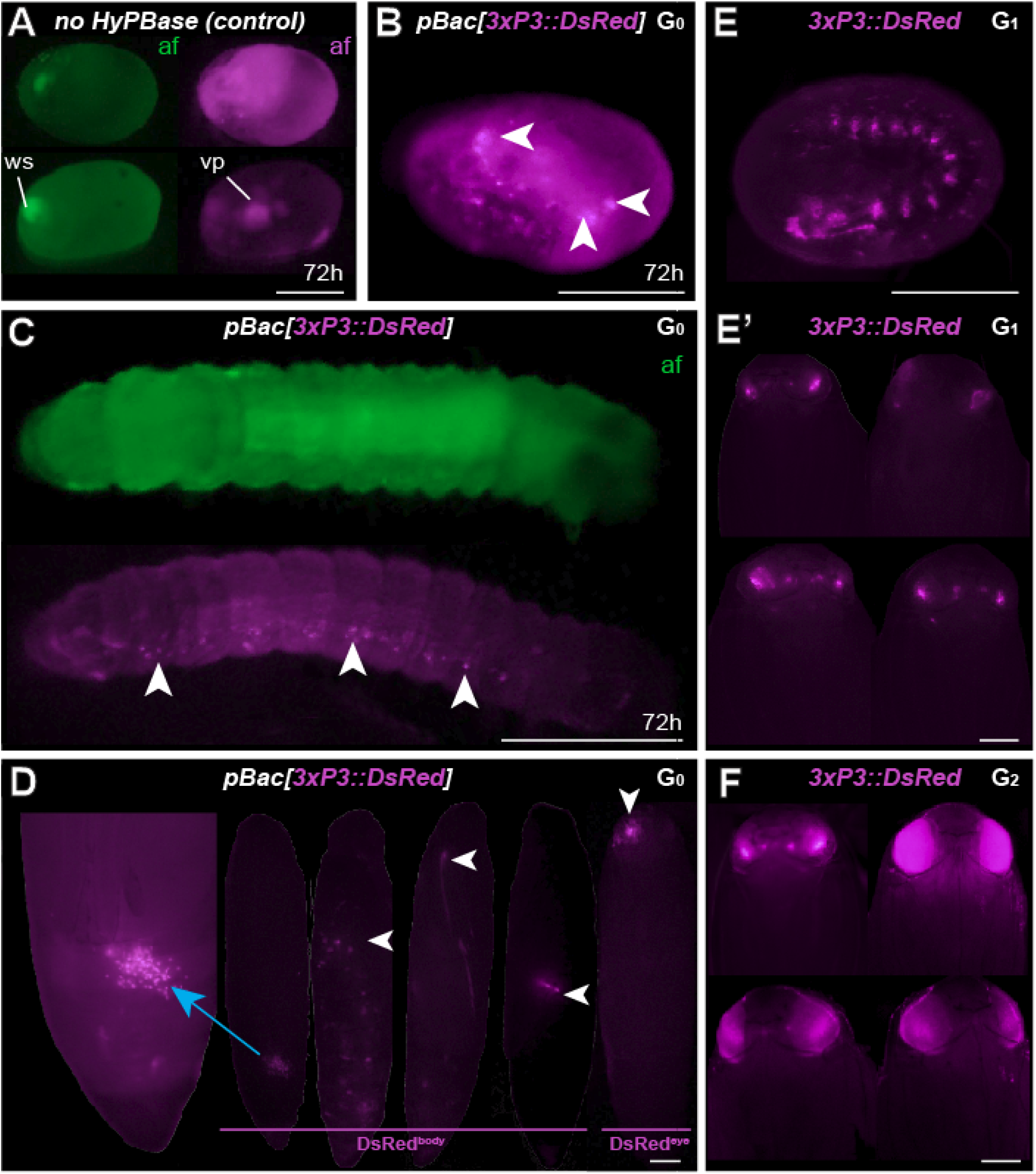
Somatic and germline transgenesis of *3xP3:DsRed*. **(A)** Two control eggs (top and bottom rows) injected with only *pBac[3xP3::DsRed]* show background autofluorescence levels in the DsRed channel (af, magenta) at 72 h post-injection, including residual signal in vitellophages (vp). Wound site (ws) autofluorescence is limited to the EGFP channel (af, green). **(B-C)** *hyPBase* mRNA and *pBac[3xP3::DsRed]* result in glial expression of *3xP3::DsRed* (magenta) in injected embryos. Ocellar expression was not observed in these experiments at the G_0_ phase. **(D)** G_0_ pupae showing various fluorescent signals in the abdomen(DsRed^body^), a phenomenon not observed with other constructs. Expression in the head (DsRed^eye^) was occasionally seen at the G_0_ phase. **(E)** G_1_ transgenic embryo with non-mosaic expression of *3xP3::DsRed*. **(E’)** G_1_ pupae showing weak eye fluorescent signals. These signals did not expand to the entire eye as the pupae developed, suggesting possible epigenetic effects. **(F)** G_2_ *Plodia* transgenic pupae obtained from G_1_ outcrosses resulted in pupae with bright *3xP3* fluorescence patterns that expanded throughout development. Variable intensity may be due to transgene copy number variation in this line. Scale bars: A-C, E = 200 μm; D, E’, F = 500 μm.

The presence of DsRed in abdominal regions suggested successful integration of the donor plasmid including in tissues close to the germline. To evaluate the germline transmission in G_0_ DsRed^eye^ individuals, we backcrossed DsRed^body^ individuals to the uninjected stock. Only 2 out of 14 G_0_ DsRed^eye^ backcrossed pairs gave G_1_ progeny (Table 2), and none inherited any DsRed fluorescence expression. In contrast, we recovered eggs from 5 G_0_ DsRed^body^ individuals that were incrossed liberally in a container, and showed full embryonic *3xP3::DsRed* signals (Fig 3E). This salvaged stock resulted in 6 G_1_ pupae with DsRed expression in the eyes (Fig. 3E’) out of 52 isolated G_1_ pupae (11.5%), with no body phenotype observed. These six G_1_ DsRed^+^ *Plodia* were then individually crossed with *Pi_wFog*, 3 of which generated 83%, 70%, and 67% G_2_ DsRed^+^ progeny (Fig. 3F). As these ratios deviate from the 1:1 ratio expected in these crosses, we conclude that more than one insert occured in the parental G_0_ founder germline.

The *3xP3* activity in this DsRed donor plasmid showed inconsistent results not seen with the EGFP donor, including absence of activity in G_0_ eye tissues, unusual abdominal fluorescent patches in G_0_ pupae, and reduced activity in G_1_ eyes. Intriguingly, full *3xP3::DsRed* activity was recovered in G_2_ pupae, suggesting possible epigenetic - effects of transient nature in earlier generations. This unusual behavior may be due to minor differences in the cassette proximal promoter (Fig. S1), to the compact design of this cassette (Fig. 1B), or to other sequence features making the insert prone to abnormal expression.

### Generation of 3xP3::EYFP transgenic lines carrying an attP docking site

We co-injected the *pBac[3XP3::EYFP; attP]* plasmid (Stern et al., 2017) with *hyPBase* mRNA into *Pi_wFog*. This donor includes an *attP docking* site (Fig. 1B), a feature that may facilitate genetic engineering using site-specific recombination, if successfully integrated into the *Plodia* genome. Control injections show little background autofluorescence and vitellophage signals under the EYFP filter set (Fig. 4A). Transgenic G_0_ embryos and larvae showed strong somatic *3xP3* activity consistent with ocular and glial expression (Fig. 4B-C), with expected mosaic variations such as unilateral expression in one side of ocellus glia and ocelli-only expression. We recovered 14 pupae with mosaic G_0_ EYFP expression (Fig. 2D) from a total of 63 surviving pupae, out of 797 embryos injected over two trials (Table 1).

**FIGURE 4.**
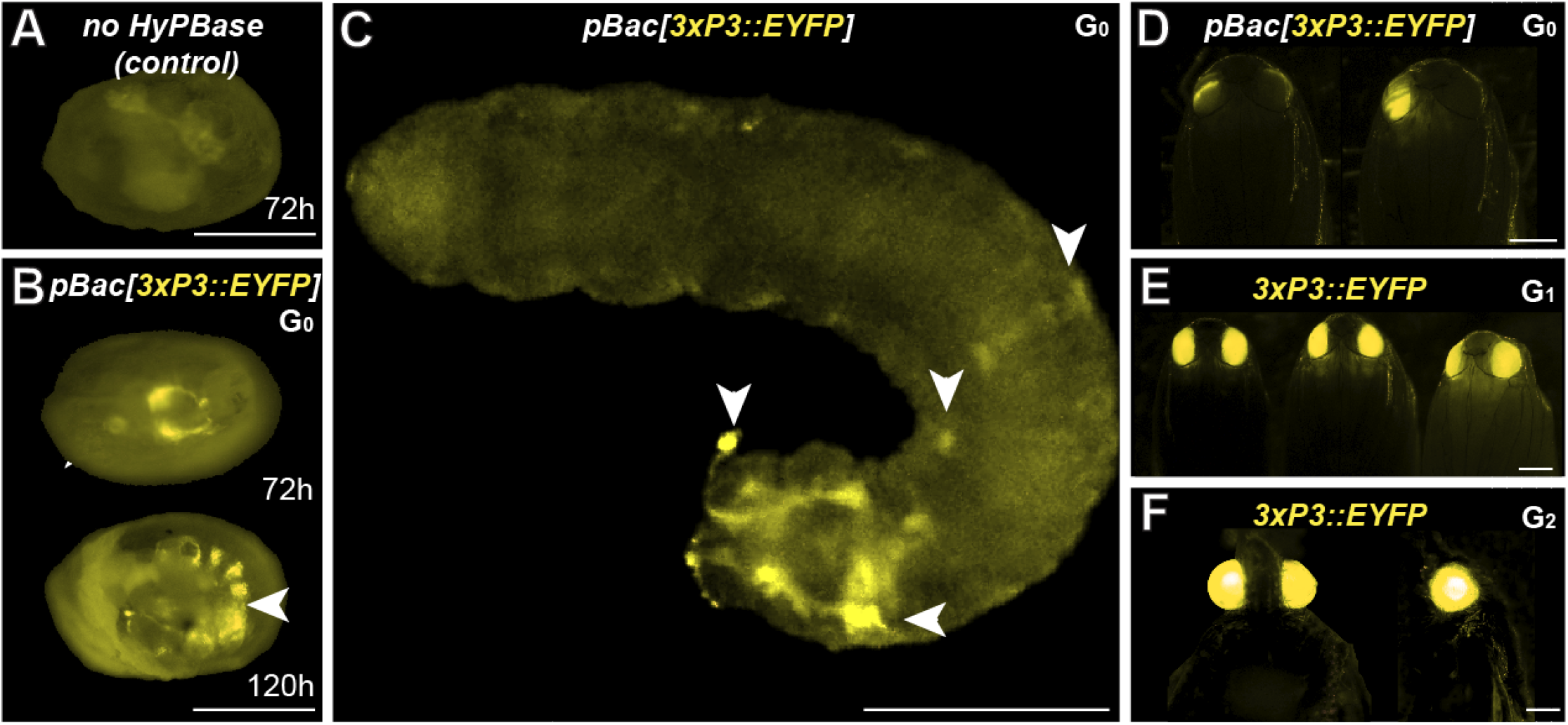
Somatic and germline transgenesis of *3xP3::EYFP* in *Plodia*. **(A)** Weak background autofluorescence in the EYFP observation channel following control injection of the donor plasmid only. **(B)** Somatic activity of *3xP3::EYFP* transgenes at 72 h and 120 h post injection in the late egg stage **(C)** Mosaic G_0_ *3xP3::EYFP* expression in a first instar larva, marking glia and ocellar stemmata (arrowheads). **(D)** Mosaic G_0_ *3xP3::EYFP* expression in pupal eyes. **(E)** *3xP3::EYFP* expression in G_1_ *Plodia* pupae. **(E)** Ventral (left) and lateral views (right) of *3xP3::EYFP* expression in G_2_ *Plodia* adults. Scale bars: A-C = 200 μm; D-F = 500 μm.

To estimate the efficiency of germline integration from these mosaic founders, we individually backcrossed the 14 EYFP^+^ G_0_ adults to single *Pi_wFog* individuals (Fig. 4E, Table 2). Seven pairs gave progeny, among which only 1 cross generated progeny with 22 G_1_ EYFP^+^ pupal phenotypes out of 37 total isolated pupae, a ratio statistically close to the 50% proportion expected from a germline tissue heterozygous for a single insertion in the G_0_ founder (^*2*^=1.32; *df* =1, *p=0*.*25*). To test if positive G_1_ individuals were heterozygous carriers for a single insertion, we simultaneously backcrossed 12 G_1_ EYFP^+^ to *Pi_wFog* and in-crossed 5 pairs of G_1_ EYFP^+^. Three of these crosses resulted in G_2_ EYFP^+^ progenies, with 59% and 48% positive ratios matching the 50% expected from backcrossing (0.18< ^*2*^< 3.17; *df* =1; 0.07< *p*< 0.67), and a 71% positive ratio matching the expected 75% in the in-cross (^*2*^= 0.84, df =1 *p* = 0.36). In summary, injection of *pBac[3XP3::EYFP; attP]* had a somatic transformation efficiency of 1.8%. The high level of mosaicism in G_0_ resulted in only 1 out of 14 successful backcrosses, resulting in a germline transformation efficiency of 0.13% (Fig. 1D, GTE = 7.1.% x 1.8% G_0_ founders), but this event was successfully carried into a stable transgenic line.

## DISCUSSION

### Transformation efficiency rates of hyPBase in Plodia

In this study, we carried out somatic and stable germline transformation in *Plodia interpunctella* using the *hyperactive piggyBac* transposase (Yusa et al., 2011), and achieved high rates of somatic transformation with 3 independent *piggyBac* donor plasmids. We injected the transposase as a mRNA and used *hyPB*^*apis*^, a version of *hyPBase* codon-optimized for honeybees (Otte et al., 2018). Because *Apis* and *Plodia* both have an average GC content of around 35% (Jørgensen et al., 2007; Kawahara et al., 2022), we can reasonably expect compatibility in their codon usage biases.

Our study is the second to use an *hyPBase* mRNA as a transposase for transgenesis in a lepidopteran insect (Chen and Palli, 2021). *Plodia* injections generated 15-40% of G_0_ somatic transformants when observed in 72-h embryos (mean of 22%), suggesting highly efficient integration.

Across different trials, a mean of 2.5% of injected eggs expressed the transgene marker as adults, representing 30% of surviving adults. However, somatic fluorescence in the injected generation does not guarantee that the transgene has transposed into the germline, or that transgenic gametes are fertile. To assess transgene inheritability into the G_1_ generation, we backcrossed G_0_ fluorescent founders to non-transgenic individuals. We obtained 3 independent G_1_ lines expressing *3xP3::EGFP* out of 6 fertile G_0_ crosses, and 1 line expressing *3xP3::EYFP* out of 7 fertile G_0_ crosses. Founders expressing *3xP3::DsRed* showed unusual patterns of G_0_ mosaicism, possibly due to epigenetic regulatory effects (see Results section), and failed to propagate the transgene when mated in single outcrossing pairs (N=14), but we recovered a stable insertion from G_1_ eggs that had been laid in a container where 5 G_0_ founders had been left to mate randomly, meaning that 1 out of 19 G_0_ transmitted *3xP3::DsRed*.

In summary, our *hyPBase* mRNA-based injections in *Plodia* resulted in germline transformation efficiency rates of 0.18% (DsRed), 0.25% (EYFP), and 0.94% (EGFP). For comparison, *Plutella* transgenic experiments using *PBase* have efficiency rates of 0.43-0.65% (Gregory et al., 2016). Our *Plodia* injection protocol has a median pharate survival of 9%, much lower than the published *Plutella* adult survival rate of 27.8% (Gregory et al., 2016). Indeed, our injection methods favor speed and quantity over precision, using relatively wide-open needle bores that avoid clogging during injections, as well as a rapid but aggressive glue-based egg sealing procedure (Heryanto et al., 2022). Only 10-25% of eggs injected with *piggyBac* reagents hatched across trials in our conditions — as opposed to 21-60% in a previous *Plodia* microinjection report conducted by another group (Bossin et al., 2007) — but this is balanced by the fact that a single experimenter can inject about 400 pre-blastoderm embryos in a 2 h session with our procedure. Overall, the germline efficiency rates reported here mean that one fertile G_0_ founder was obtained for every 106 (EGFP), 555 (DsRed), and 777 (EYFP) injected embryos, making a 2-4 h injection effort (400-800 eggs) reasonably well suited for initiating each transgenic line attempt. Ultimately, practicality boils down to a trade-off between the number of injected embryos and their survival, and our data suggest that the high efficiency of *hyPBase* (Yusa et al., 2011; Eckermann et al., 2018) can make transgenesis feasible if one of these two factors is not optimal.

### Other practical considerations for transgenesis in Lepidoptera

Mendelian segregation patterns observed at the G_2_ generations indicate that all 4 out of 5 stable lines originated as single-insertion events, with G_0_ founders likely carrying a single copy (Table 2). This feature can be used by experimenters to use various crossing strategies in the future, but we must highlight that single-mating strategies and crossing conditions resulted in few successful pairings in our initial attempts (*e*.*g*. 11-50% of G_0_ crosses, Table 2). This artificially lowered germline transmission rates, likely due to founders failing to mate in small containers in suboptimal condition. As we gained experience with *Plodia* husbandry during these experiments, we increased mating success rates to 66-78% in subsequent generations (see Methods for the optimized procedure). Furthermore we recommend to mix one transgene carrier with 2-3 wild-type unmated adults of the opposite sex instead of one, as this maximizes the likelihood of successful mating in this system (Brower, 1975; Huang and Subramanyam, 2003).

Each of the three constructs we tested provided complementary information. The EGFP construct was the largest and delivered the highest germline transformation rate. Of note, the compact DsRed construct resulted in unusual G_0_ fluorescent patterns. While circumstantial, these observations bode well for using large inserts, and we caution that the *pBac[3xP3::DsRed]* (*pHD-DsRed*, Addgene #64703) has a more compact minimal promoter that might also explain its weaker expression (Fig. S1). The candidate germline driver of *mScarlet*, consisting of the proximal promoter and 3’UTR of *nanos-O* (Nakao and Takasu, 2019; Xu et al., 2019) cloned from the *Plodia* genome, failed to drive detectable fluorescence in ovarian tissues. We will investigate alternative germline promoters in the future (Xu et al., 2022), for instance by testing the *PhiC31* site-specific integrase at the *attP* docking site from our new EYFP transgenic line (Yonemura et al., 2013; Haghighat-Khah et al., 2015; Stern et al., 2017; Stern, 2022). Both EGFP and EYFP showed robust and strong *3xP3*-driven expression at all generations, without noticeable decrease over time in adult eyes (Das Gupta et al., 2015), with EYFP benefiting from lower autofluorescence effects than EGFP at various stages. We strategically used a *white* mutant strain deficient for eye-screening pigment, as routinely done in other insects to facilitate the screening of *3xP3*-driven fluorescence (Stern et al., 2017; Klingler and Bucher, 2022), and this mutation also increases the translucency of *Plodia* larvae (Shirk, 2021; Heryanto et al., 2022). Of note, *white* mutations can be recessive-lethal in some lepidopteran species (Khan et al., 2017). Until alternative way to generate depigmented eyes are found, this may limit the usefulness of *3xP3* drivers, especially in species where eggs and larvae are opaque and where screening becomes limited to narrow developmental windows (Das Gupta et al., 2015; Özsu et al., 2017). In such species, we suggest that stronger, more ubiquitous promoters of viral origin such as *Op-ie2* and *Hr5-ie1* may be more practical for transgenic screening (Martins et al., 2012; Xu et al., 2019, 2022).

## MATERIALS AND METHODS

### Plodia strains and rearing

The *Pi_wFog* strain (Heryanto et al., 2022) consists of an introgression of the recessive *w-* mutation (Shirk, 2021) from the *Pi* ^*w-*^ strain (origin: USA, kind gift of Paul Shirk), into the genetic background of the “Dundee” strain (origin : UK, kind gift of Mike Boots). Genome assemblies of both *Pi* ^*w-*^ and *Pi_Dundee* parental strains are available (Roberts et al., 2020; Kawahara et al., 2022). The resulting hybrid *Pi_wFog* strain has been maintained in inbred state for 3 years and used throughout this study. All rearing used previously published methods (Heryanto et al., 2022), using special containers and a wheat bran-sucrose-glycerol diet (Silhacek and Miller, 1972). A rearing temperature of 28°C resulted in a generation time of 28 d.

### Plasmid constructs

The *pBac[3xP3::EGFP; Tc’hsp5’-Gal4Delta-3’UTR]* (Addgene plasmid # 86449) was used as a donor plasmid with Piggybac insertion repeats and the 3xP3::EGFP reporter (Schinko et al., 2010). To generate *pBac[3xP3::EGFP; nosO_prom::mScarlet-nosO_3’UTR]*, an *mScarlet* cassette preceded by 2 kb of 5’UTR sequence immediately upstream of the *Plodia nanos-O* start codon, was synthesized in the *pUC-GW-Amp* backbone by Genewiz (South Plainfield, NJ) and sub-cloned into the *FseI* and *AscI* restriction sites of *pBac[3xP3::EGFP; Tc’hsp5’-Gal4Delta-3’UTR]*. The *pBac[3xP3::DsRed]* (*pHD-DsRed*) and *pBac[3XP3::EYFP; attP]* plasmids were obtained from Addgene (#64703, and #86860) and used without modification (Gratz et al., 2014, 2015; Stern et al., 2017). All the *3xP3-*driven fluorophore genes included an *SV40* termination sequence.

### Transposase mRNA and injection mixes

The *pGEM-T_hyPB*^*apis*^ plasmid encodes a *hyPBase* that was codon-optimized for honeybees (Otte et al., 2018). The source plasmid was purified using the QIAprep Spin Miniprep Kit (Qiagen, Germantown, MD), linearized with *Nco*I-HF (NEB, Ipswich, MA) and concentrated using acetate/ethanol precipitation. Around 500 ng of linearized template were transcribed using the Invitrogen™ mMESSAGE mMACHINE™ T7 ULTRA Transcription Kit (Invitrogen, Carlsbad, CA) and purified using the MEGAclear™ Transcription Clean-Up Kit (Invitrogen, Carlsbad, CA). After quantification with Nanodrop (Thermofisher, Waltham, MA), the solution was divided into 1050 ng/μL one-time use aliquots and stored at -80°C.

### Microinjections

Injection mixes consisted of 400 ng/μl *hyPBase* mRNA, 200 ng/μl donor plasmid, and 0.05% cell-culture grade Phenol Red (Sigma-Aldrich, Burlington, MA). Donor plasmids without *hyPBase* mRNA were injected in separate experiments as controls. Microinjection procedures (Figure 1A) followed a previously described procedure (Heryanto et al., 2022), with all embryo injections performed within 40 min after egg laying (AEL). Injected embryos were counted and kept in a rearing container with a small damp Kimwipe at 28°C. For the first 72 h, the container vent was covered with tape in order to maintain humidity saturation, a parameter that prevents egg desiccation. After 72 h, the vent was opened and the Kimwipe removed, and about five flakes of *Plodia* food added next to the eggs, in order to keep the emerging larvae within the injection dish. Mean emergence time of the *Pi_wFog* strain is 83 h AEL at 28°C for uninjected eggs, and is delayed by injection stress to 100-115 h AEL. Because of this variability, we report times of observation after injection in hours rather than in relative percentages.

### Fluorescent microscopy

Larvae and adult *Plodia* were anesthetized in tissue culture dishes positioned over a cold metal block during microscopy observation. All pictures were taken under the Olympus SZX16 stereomicroscope equipped with a Lumencor SOLA Light Engine SM 5-LCR-VA lightsource or standard stereomicroscope brightfield lamp, and with a trinocular tube connected to an Olympus DP73 digital color camera. Separation of fluorescent channels was performed using Chroma Technology filter sets ET-EGFP 470/40x 510/20m, ET-EYFP 500/20x 535/30m, and AT-TRICT-REDSHFT 540/25x, 620/60m.

### Survival and G_0_ somatic transformation rates

Embryonic survival rates (“egg hatching” rates) were determined by the ratio of hatched eggs at 120 h AEL over the number of injected eggs (*N*_*inj*_). Empty egg shells were counted for this purpose instead of first-instar hatchlings, which are difficult to count accurately in the presence of food. Pharate survival rates were determined by the ratio of pupae obtained from a given injection experiment, divided by *N*_*inj*_, and thus accounts for mortality occurring at embryonic and larval stages. Pupal mortality was negligible, making pharate survival rates a reasonable proxy for overall adult survival, and is more convenient to couple to fluorescent screening than in mobile adults. G_0_ transformation rates were independently measured in embryos and in pupae. For embryos, eggs with bright, internal fluorescent signals consistent with an ocellar or glial expression were counted as positive (fluorescent, F+ in Table 1) around 72 h AEL, and non-fluorescent eggs were counted as negative (F-). To isolate individual pupae, cardboard strips that are preferentially used as pupation sites (“hotels”) were added into containers containing fifth instar larvae, allowing a convenient isolation of individual *Plodia* pupae. Pupae were then extirpated from these lodges and aligned on double-sided tape for fluorescence screening. Pupae with any glial or eye signal were counted as positive, while others were counted as negative. G_0_ somatic transformation efficiency rate was determined as the number of healthy adult individuals emerged from fluorescent pupae, and normalized by *N*_*inj*_.

### Controlled crosses for germline transmission

Germline transformation efficiency rates factored the somatic transformation efficiency rate by the proportion of attempted G_0_ backcrosses yielding transgenic offspring. G_0_ transgenic adults or late pupae exhibiting positive fluorescent signals (G_0_ F+) were crossed to a single unmated *Pi_wFog* adult of the opposite sex, by mixing in a 1.25 oz Plastic Souffle Cup (Solo) containing ∼ 0.2 g of diet and ∼ 1 cm^2^ of paper towel. *Pi_wFog* outcrossing mates were replaced if found dead before any visible egg laying. These cups were monitored for up to 2 weeks for any larval emergence, after which they were transferred to a vented rearing container with a bed of *Plodia* food (modified LocknLock containers described in Heryanto *et al*., 2022; 177 mL and 350 mL formats). At the wandering L5 stage, cardboard “hotels” were added into the containers for pupal isolation. G_1_ pupae with positive fluorescent signal were counted and backcrossed to an unmated *Pi_wFog* with the same procedure stated above. The resulting G_2_ pupae were in-crossed as sib-matings and maintained as isogenic stock in the G_3_ generations and henceforth.

## DATA AVAILABILITY STATEMENT

The *pBac[3xP3::EGFP; nosO_prom::mScarlet-nosO_3’UTR]* plasmid generated in this study is available upon request.

## AUTHOR CONTRIBUTIONS

CH and AM designed the study and wrote the manuscript. AMV and AM advised on the methodology. CH performed the experiments and analyzed the data.

## SUPPLEMENTARY MATERIAL

The Supplementary Material for this article can be found online at: http://____

**FIGURE S1.**
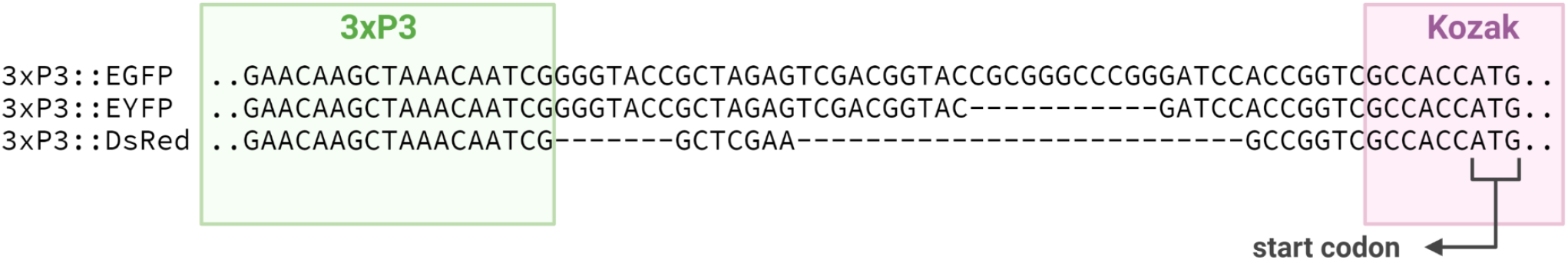
Alignment of the region spanning, from left to right, the 3’ end of the *3xP3* promoter and Kozak sequence preceding the fluorophore genes in 3 *pBac* donor plasmids.

## REFERENCES

Bartlett, L. J., Wilfert, L., and Boots, M. (2018). A genotypic trade-off between constitutive resistance to viral infection and host growth rate. Evolution 72, 2749–2757. doi: 10.1111/evo.13623.

Berghammer, A. J., Klingler, M., and A Wimmer, E. (1999). A universal marker for transgenic insects. Nature 402, 370–371.

Bossin, H., Furlong, R. B., Gillett, J. L., Bergoin, M., and Shirk, P. D. (2007). Somatic transformation efficiencies and expression patterns using the JcDNV and piggyBac transposon gene vectors in insects. Insect Mol. Biol. 16, 37–47. doi: 10.1111/j.1365-2583.2006.00693.x.

Brower, J. H. (1975). Plodia interpunctella: effect of sex ratio on reproductivity. Ann. Entomol. Soc. Am. 68, 847–851.

Chen, X., and Palli, S. R. (2021). Hyperactive piggyBac Transposase-mediated Germline Transformation in the Fall Armyworm, Spodoptera frugiperda. JoVE J. Vis. Exp., e62714.

Das Gupta, M., Chan, S. K. S., and Monteiro, A. (2015). Natural loss of eyeless/Pax6 expression in eyes of Bicyclus anynana adult butterflies likely leads to exponential decrease of eye fluorescence in transgenics. PloS One 10, e0132882.

Dyby, S., and Silhacek, D. L. (1997). Juvenile hormone agonists cause abnormal midgut closure and other defects in the moth, Plodia interpunctella (Lepidoptera: Pyralidae). Invertebr. Reprod. Dev. 32, 231–244.

Eckermann, K. N., Ahmed, H. M. M., KaramiNejadRanjbar, M., Dippel, S., Ogaugwu, C. E., Kitzmann, P., et al. (2018). Hyperactive piggyBac transposase improves transformation efficiency in diverse insect species. Insect Biochem. Mol. Biol. 98, 16–24. doi: 10.1016/j.ibmb.2018.04.001.

Gratz, S. J., Rubinstein, C. D., Harrison, M. M., Wildonger, J., and O’Connor-Giles, K. M. (2015). CRISPR-Cas9 genome editing in Drosophila. Curr. Protoc. Mol. Biol. 111, 31–2.

Gratz, S. J., Ukken, F. P., Rubinstein, C. D., Thiede, G., Donohue, L. K., Cummings, A. M., et al. (2014). Highly specific and efficient CRISPR/Cas9-catalyzed homology-directed repair in Drosophila. Genetics 196, 961–971.

Gregory, M., Alphey, L., Morrison, N. I., and Shimeld, S. M. (2016). Insect transformation with piggyBac: getting the number of injections just right. Insect Mol. Biol. 25, 259–271.

Haghighat-Khah, R. E., Scaife, S., Martins, S., St John, O., Matzen, K. J., Morrison, N., et al. (2015). Site-Specific Cassette Exchange Systems in the Aedes aegypti Mosquito and the Plutella xylostella Moth. PLOS ONE 10, e0121097. doi: 10.1371/journal.pone.0121097.

Handler, A. M. (2002). Use of the piggyBac transposon for germ-line transformation of insects. Insect Biochem. Mol. Biol. 32, 1211–1220.

Harrison, P. W., Mank, J. E., and Wedell, N. (2012). Incomplete Sex Chromosome Dosage Compensation in the Indian Meal Moth, Plodia interpunctella, Based on De Novo Transcriptome Assembly. Genome Biol. Evol. 4, 1118–1126. doi: 10.1093/gbe/evs086.

Heryanto, C., Hanly, J. J., Mazo-Vargas, A., Tendolkar, A., and Martin, A. (2022). Mapping and CRISPR homology-directed repair of a recessive white eye mutation in Plodia moths. iScience 25, 103885. doi: 10.1016/j.isci.2022.103885.

Horn, C., Schmid, B. G. M., Pogoda, F. S., and Wimmer, E. A. (2002). Fluorescent transformation markers for insect transgenesis. Insect Biochem. Mol. Biol. 32, 1221–1235. doi: 10.1016/S0965-1748(02)00085-1.

Huang, F., and Subramanyam, B. (2003). Effects of delayed mating on reproductive performance of Plodia interpunctella (Hübner)(Lepidoptera: Pyralidae). J. Stored Prod. Res. 39, 53–63.

Jørgensen, F. G., Schierup, M. H., and Clark, A. G. (2007). Heterogeneity in regional GC content and differential usage of codons and amino acids in GC-poor and GC-rich regions of the genome of Apis mellifera. Mol. Biol. Evol. 24, 611–619.

Kawahara, A. Y., Storer, C. G., Markee, A., Heckenhauer, J., Powell, A., Plotkin, D., et al. (2022). Long-read HiFi sequencing correctly assembles repetitive heavy fibroin silk genes in new moth and caddisfly genomes. Gigabyte 2022, 1–14. doi: 10.46471/gigabyte.64.

Khan, S. A., Reichelt, M., and Heckel, D. G. (2017). Functional analysis of the ABCs of eye color in Helicoverpa armigera with CRISPR/Cas9-induced mutations. Sci. Rep. 7, 1–14.

Klingler, M., and Bucher, G. (2022). The red flour beetle T. castaneum: elaborate genetic toolkit and unbiased large scale RNAi screening to study insect biology and evolution. EvoDevo 13, 1–11.

Kristensen, N. P., Scoble, M. J., and Karsholt, O. L. E. (2007). Lepidoptera phylogeny and systematics: the state of inventorying moth and butterfly diversity. Zootaxa 1668, 699–747.

Laptev, I. A., Raevskaya, N. M., Filimonova, N. A., and Sineoky, S. P. (2017). The piggyBac Transposon as a Tool in Genetic Engineering. Appl. Biochem. Microbiol. 53, 874–881.

Martins, S., Naish, N., Walker, A. S., Morrison, N. I., Scaife, S., Fu, G., et al. (2012). Germline transformation of the diamondback moth, Plutella xylostella L., using the piggyBac transposable element: Germline transformation of diamondback moth. Insect Mol. Biol. 21, 414–421. doi: 10.1111/j.1365-2583.2012.01146.x.

Mbata, G. N. (1985). Some physical and biological factors affecting oviposition by Plodia interpunctella (Hubner) (Lepidoptera: Phycitidae). Int. J. Trop. Insect Sci. 6, 597–604. doi: 10.1017/S1742758400009176.

Nakao, H., and Takasu, Y. (2019). Complexities in Bombyx germ cell formation process revealed by Bm-nosO (a Bombyx homolog of nanos) knockout. Dev. Biol. 445, 29–36. doi: 10.1016/j.ydbio.2018.10.012.

Otte, M., Netschitailo, O., Kaftanoglu, O., Wang Y. Jr, R. E. P., and Beye, M. (2018). Improving genetic transformation rates in honeybees. Sci. Rep. 8, 16534. doi: 10.1038/s41598-018-34724-w.

Özsu, N., Chan, Q. Y., Chen, B., Gupta, M. D., and Monteiro, A. (2017). Wingless is a positive regulator of eyespot color patterns in Bicyclus anynana butterflies. Dev. Biol. 429, 177–185.

Roberts, K. E., Meaden, S., Sharpe, S., Kay, S., Doyle, T., Wilson, D., et al. (2020). Resource quality determines the evolution of resistance and its genetic basis. Mol. Ecol. 29, 4128–4142. doi: https://doi.org/10.1111/mec.15621.

Roskov, Y., Kunze, T., Paglinawan, L., Orrell, T., Nicolson, D., Culham, A., et al. (2013). Species 2000 & ITIS Catalogue of Life, 2013 Annual Checklist.

Schinko, J. B., Weber, M., Viktorinova, I., Kiupakis, A., Averof, M., Klingler, M., et al. (2010). Functionality of the GAL4/UAS system in Tribolium requires the use of endogenous core promoters. BMC Dev. Biol. 10, 1–12.

Shirk, B. D. (2021). Gene Editing of the ABC Transporter/White Locus Using CRISPR/Cas9 Mutagenesis in the Indian meal moth (Plodia interpunctella).

Silhacek, D. L., and Miller, G. L. (1972). Growth and Development of the Indian Meal Moth, Plodia interpunctella (Lepidoptera: Phycitidae), Under Laboratory Mass-Rearing Conditions1. Ann. Entomol. Soc. Am. 65, 1084–1087. doi: 10.1093/aesa/65.5.1084.

Stern, D. L. (2022). Transgenic tools for targeted chromosome rearrangements allow construction of balancer chromosomes in non-melanogaster Drosophila species. G3 12, jkac030.

Stern, D. L., Crocker, J., Ding, Y., Frankel, N., Kappes, G., Kim, E., et al. (2017). Genetic and transgenic reagents for Drosophila simulans, D. mauritiana, D. yakuba, D. santomea, and D. virilis. G3 Genes Genomes Genet. 7, 1339–1347.

Tang, P.-A., Wu, H.-J., Xue, H., Ju, X.-R., Song, W., Zhang, Q.-L., et al. (2017). Characterization of transcriptome in the Indian meal moth Plodia interpunctella (Lepidoptera: Pyralidae) and gene expression analysis during developmental stages. Gene 622, 29–41. doi: 10.1016/j.gene.2017.04.018.

Thomas, J. L., Da Rocha, M., Besse, A., Mauchamp, B., and Chavancy, G. (2002). 3xP3-EGFP marker facilitates screening for transgenic silkworm Bombyx mori L. from the embryonic stage onwards. Insect Biochem. Mol. Biol. 32, 247–253.

Xu, J., Chen, R., Chen, S., Chen, K., Tang, L., Yang, D., et al. (2019). Identification of a germline-expression promoter for genome editing in Bombyx mori. Insect Sci. 26, 991–999. doi: 10.1111/1744-7917.12657.

Xu, X., Harvey-Samuel, T., Siddiqui, H. A., Ang, J. X. D., Anderson, M. E., Reitmayer, C. M., et al. (2022). Toward a CRISPR-Cas9-based gene drive in the diamondback moth Plutella xylostella. CRISPR J. 5, 224–236.

Yonemura, N., Tamura, T., Uchino, K., Kobayashi, I., Tatematsu, K., Iizuka, T., et al. (2013). phiC31-integrase-mediated, site-specific integration of transgenes in the silkworm, Bombyx mori (Lepidoptera: Bombycidae). Appl. Entomol. Zool. 48, 265–273.

Yusa, K., Zhou, L., Li, M. A., Bradley, A., and Craig, N. L. (2011). A hyperactive piggyBac transposase for mammalian applications. Proc. Natl. Acad. Sci. 108, 1531–1536.

